# Humoral signaling-mediated effects of unilateral brain injury: differences in the left-right sided afferent responses

**DOI:** 10.1101/2022.04.15.488460

**Authors:** Hiroyuki Watanabe, Yaromir Kobikov, Daniil Sarkisyan, Igor Lavrov, Jens Schouenborg, Mengliang Zhang, Georgy Bakalkin

## Abstract

Disruption of neural tracts descending from the brain to the spinal cord after brain trauma and stroke causes postural and sensorimotor deficits. We previously showed that unilateral lesion to the sensorimotor cortex in rats with completely transected thoracic spinal cord produced asymmetry in hindlimb posture and withdrawal reflexes. Supraspinal signals to hindlimb muscles may be transmitted through the paravertebral chain of sympathetic ganglia that remain intact after the transection. We here demonstrated that prior transection of the spinal cord at the cervical level that was rostrally to segments with preganglionic sympathetic neurons, did not abolish formation of asymmetry in hindlimb posture and musculo-articular resistance to stretch after unilateral brain injury. Thus not the sympathetic system but humoral signals may mediate the effects of brain injury on the lumbar spinal circuits. The asymmetric responses in rats with transected spinal cords were eliminated by bilateral lumbar dorsal rhizotomy after the left-side brain injury, but resistant to deafferentation after the right-side brain lesion. Two mechanisms, one dependent on and one independent of afferent input may account for asymmetric hindlimb motor responses. Resistance to deafferentation may be due to sustained stretch- and effort-unrelated muscle contractions that is often observed in patients with central lesions. Left-right asymmetry is unusual feature of these mechanisms that both are activated by humoral signals.

**Graphical Abstract:** 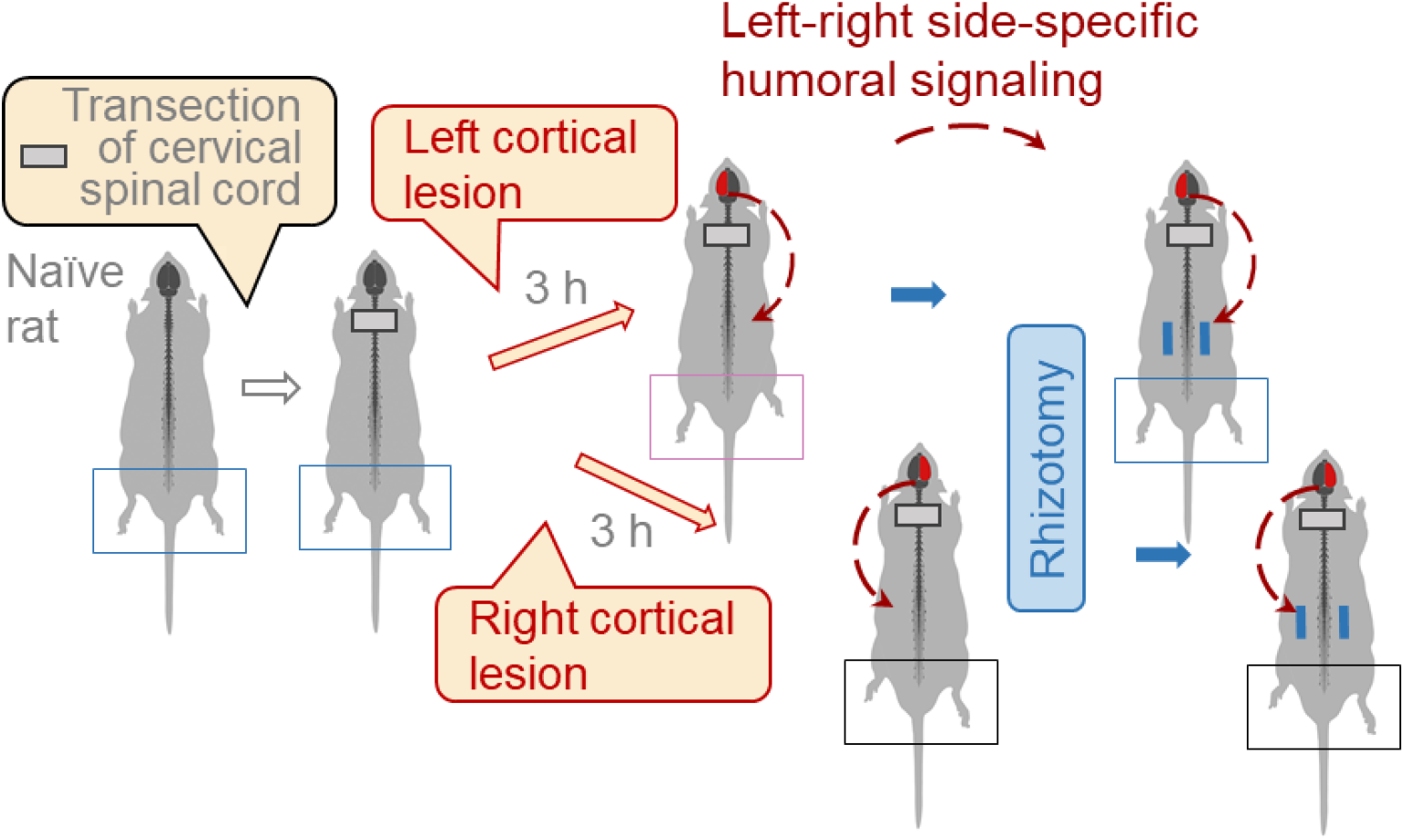

## Introduction

External symmetry and functional specialization of the left and right hemispheres are fundamental principles of the central nervous system organization (Concha et al., 2012; Duboc et al., 2015; Gunturkun and Ocklenburg, 2017; MacNeilage et al., 2009). Language, emotions, and spatial cognition are lateralized. Asymmetries of neural circuits subserving these functions may be a basis for the lateralization (Concha et al., 2012; Horstick et al., 2020). Several lines of evidence demonstrate that the lateralized neurocircuits are regulated by neurohormones that operate either on the left or right side (Allen et al., 2021; Deliagina et al., 2000; Hussain et al., 2012; Kawakami et al., 2003; Kononenko et al., 2017; Kononenko et al., 2018; Marlin et al., 2015; Nation et al., 2018; Phelps et al., 2019; Watanabe et al., 2015; Zink et al., 2011). Thus, oxytocin enhances responses of the left but not right auditory cortex that enables retrieval behavior (Marlin et al., 2015). The dynorphin – κ-opioid and CGRP neurocircuits control lateralized processing of nociceptive information in the amygdala (Allen et al., 2021; Nation et al., 2018; Phelps et al., 2019). Intriguing questions are if this neurohormonal mechanism is bipartite and differentially controls mirror symmetric neurocircuits on their left and right sides, and if it operates locally in the CNS or may span the entire body by virtue of neurohormones released from the hypothalamus and pituitary gland.

Traumatic brain injury and stroke in patients, and brain lesions in animal experiments cause postural and sensorimotor deficits that are generally contralesional and include asymmetric posture and reflexes (Dewald et al., 1999; Serrao et al., 2012; Spaich et al., 2006; Wilson et al., 2017; Zhang et al., 2020). After a unilateral injury to the hindlimb sensorimotor cortex, animals exhibit the hindlimb postural asymmetry (HL-PA) with contralesional limb flexion, differences in musculo-articular resistance of the contra- and ipsilesional hindlimbs to stretch, and asymmetry of the nociceptive withdrawal reflexes (NWRs) with greater activity on the contra-*vs*. ipsilesional side (Wolpaw, 2012; Zhang et al., 2020).

These effects are induced through descending neural pathways that convey motor commands from the cortex to spinal motoneurons (Lemon, 2008; Purves, 2001; Smith et al., 2017). Early studies suggested that in addition to the descending neural tracts the contralesional effects of brain injury may be mediated by an extra-spinal mechanism (Cope et al., 1980; Wolpaw and Lee, 1989). To test this hypothesis, the descending neural tracts were disabled by complete transection of the thoracic spinal cord before the brain injury was performed (Lukoyanov et al., 2021). In spite of prior spinal cord transection, a unilateral injury of the hindlimb sensorimotor cortex resulted in formation of HL-PA with contralateral flexion, and asymmetric changes in hindlimb withdrawal reflexes and expression patterns of neuroplasticity-related genes in the lumbar spinal cord.

Hypothetically, signals from the injured brain may also reach the lumbar segments through the sympathetic system or humoral pathway. Signals may be processed through preganglionic sympathetic neurons that are located in the T1 - L2-3 spinal segments and project to prevertebral sympathetic ganglia, and then through the chain of these ganglia to the lumbar segments. To assess if the sympathetic system mediates the brain injury signals, we here analyzed the UBI effects on hindlimb posture and sensorimotor functions in rats with prior transection of the spinal cord at the cervical level that was rostrally to the spinal segments with preganglionic sympathetic neurons. We next examined if afferent input is required for development of the UBI - induced hindlimb responses through the humoral pathway. For this purpose, the spinal cord segments from L1 to S2 in rats with completely transected cervical spinal cord were bilaterally deafferentated, and the effects of rhizotomy on the UBI-induced HL-PA were analyzed. Bilateral Deafferentation was performed on the both sides to exclude the effects of afferent signals from both hindlimbs on the postural and motor output. In general, this study aims to add to understanding of regulation of the left and right sided processes in bilaterally symmetric animals through the humoral pathway that may enable transmission of the left-right side specific signals targeting the afferent and efferent mechanisms.

## Materials and methods

### Animals

Adult male Sprague Dawley rats (Taconic, Denmark) weighing 200-430 g were used in the study. The animals received food and water ad libitum, and were kept in a 12-h day-night cycle (light on from 10:00 a.m. to 10:00 p.m.) at a constant environmental temperature of 21°C (humidity: 65%) and randomly assigned to their respective experimental groups. Approval for animal experiments was obtained from the Malmö/Lund ethical committee on animal experiments (No.: M7-16. Experiments were performed from 9:00 a.m. to approximately 8:00 p.m. After the experiments were completed, the animals were given a lethal dose of pentobarbital.

### Spinal cord transection

The animals were anesthetized with sodium pentobarbital (intraperitoneal, I.P.; 60 mg/kg body weight, as an initial dose and then 6 mg/kg every hour), or with isoflurane (1.5% isoflurane in a mixture of 65% nitrous oxide and 35% oxygen) anesthesia. Core temperature of the animals was controlled using a feedback-regulated heating system.

The experimental design included rats with UBI which was preceded by a complete spinal cord transection. Anaesthetized animals were first placed on a surgery platform and the skin of the back was incised along the midline at the level of the superior thoracic vertebrae. After the back muscles were retracted to the sides, a laminectomy was performed at the C6 and C7 vertebrae. A 3-4-mm spinal cord segment between the two vertebrae was dissected and removed (Lukoyanov et al., 2021). The completeness of the transection was confirmed by (i) inspecting the cord during the operation to ensure that no spared fibers bridged the transection site and that the rostral and caudal stumps of the spinal cord were completely retracted; and (ii) examining the spinal cord in all animals after termination of the experiment.

### Brain surgery and histology

Following the transection, the rats were mounted onto the stereotaxic frame and the head was fixed in a position in which the bregma and lambda were located at the same horizontal level. After local injection of lidocaine (Xylocaine, 3.5 mg/ml) with adrenaline (2.2 μg/ml), the scalp was cut open and a piece of the parietal bone located 0.5 – 4.0 mm posterior to the bregma and 1.8 – 3.8 mm lateral to the midline (Paxinos and Watson, 2007) was removed. The part of the cerebral cortex located below the opening that includes the hind-limb representation area of the sensorimotor cortex (HL-SMC) was aspirated with a metallic pipette (tip diameter 0.5 mm) connected to an electrical suction machine (Craft Duo-Vec Suction unit, Rocket Medical Plc, UK). Care was taken to avoid damaging the white matter below the cortex. After the ablation, bleeding was stopped with a piece of Spongostone and the bone opening was covered with a piece of TissuDura (Baxter, Germany). For sham operations, animals underwent the same anesthesia and surgical procedures, but the cortex was not ablated.

After completion of all surgical procedures, the wounds were closed with the 3-0 suture (AgnTho’s, Sweden) and the rat was kept under an infrared radiation lamp to maintain body temperature during monitoring of postural asymmetry (up to 3 h) and during stretching force analysis.

Localization and size of cortical lesions were analyzed in rats with left side (n = 10) and right side (n = 11) UBI. After perfusion with 4% paraformaldehyde the brain was removed and postfixed in the same fixative overnight. Then the brain was soaked in phosphate-buffered saline with 30% sucrose for 48 hours, dissected into blocks which were then sliced into 50 µm sections with a freezing microtome. Every fourth section was stained with toluidine (Nissl stain), and all the stained sections across the lesion site were photographed and the rostrocaudal respective mediolateral extension as well as lesion volume were calculated.

### Bilateral deafferentation surgery

A subset of rats underwent bilateral deafferentation by rhizotomy 3 h after the brain surgery. A laminectomy was performed from T11 to L3 vertebral levels, then the dura was open and the spinal dorsal roots from L2 to S2 levels were cut bilaterally with a pair of fine scissors.

### Analysis of hindlimb postural asymmetry (HL-PA) by the hands-on and hands-off methods

The HL-PA value and the side of the flexed limb were assessed as described elsewhere (Lukoyanov et al., 2021; Watanabe et al., 2020; Zhang et al., 2020). Briefly, the measurements were performed under pentobarbital (60 mg/kg, i.p.). The level of anesthesia was characterized by a barely perceptible corneal reflex and a lack of overall muscle tone. The anesthetized rat was placed in the prone position on the 1-mm grid paper.

In the hands-on analysis, the hip and knee joints were straightened by gently pulling the hindlimbs backwards for 1 cm to reach the same level. Then, the hindlimbs were set free and the magnitude of postural asymmetry was measured in millimeters as the length of the projection of the line connecting symmetric hindlimb distal points (digits 2-4) on the longitudinal axis of the rat. The procedure was repeated six times in immediate succession, and the mean HL-PA value for a given rat was used in statistical analyses.

In the hands-off method, silk threads were glued to the nails of the middle three toes of each hindlimb, and their other ends were tied to one of two hooks attached to the movable platform that was operated by a micromanipulator constructed in the laboratory (Lukoyanov et al., 2021). To reduce potential friction between the hindlimbs and the surface with changes in their position during stretching and after releasing them, the bench under the rat was covered with plastic sheet and the movable platform was raised up to form a 10° angle between the threads and the bench surface. Positions of the limbs were adjusted to the same, symmetric level, and stretching was performed for the 1.5 cm distance at a rate of 2 cm/sec. The threads then were relaxed, the limbs were released and the resulting HL-PA was photographed. The procedure was repeated six times in succession, and the mean value of postural asymmetry for a given rat was calculated and used in statistical analyses.

The limb displacing shorter projection was considered as flexed. The HL-PA was measured in mm with negative and positive HL-PA values that are assigned to rats with the left and right hindlimb flexion, respectively. This measure, the postural asymmetry size (PAS) shows the HL-PA value and flexion side. The PAS does not show the proportion of the animals with asymmetry in each group, whether all or a small fraction of animals display the asymmetry; and cannot be used for analysis of rat groups with the similar number of left or right flexion. In the latter case the HL-PA value would be about zero. Therefore, the HL-PA was also assessed by the magnitude of postural asymmetry (MPA) that shows absolute flexion size, and the probability of postural asymmetry (P_A_) that shows the proportion of animals exhibiting HL-PA at the imposed threshold (> 1 mm). The P_A_ does not show flexion size and size. These three measures are obviously dependent; however, they are not redundant and for this reason, all are required for data presentation and characterization of the HL-PA.

### Analysis of hindlimb resistance to stretch

Stretching force was analyzed under pentobarbital anesthesia within 3-5 hrs after UBI using the micromanipulator-controlled force meter device constructed in the laboratory (Zhang et al., 2020). Two Mark-10 digital force gauges (model M5-05, Mark-10 Corporation, USA) with a force resolution 50 mg were fixed on a movable platform operated by a micromanipulator. Three 3-0 silk threads were glued to the nails of the middle three toes of each hindlimb, and their another ends were hooked to one of two force gauges. The flexed leg of the rat in prone position was manually stretched to the level of the extended leg; this position was taken as 0 mm point. Then both hind limbs were stretched in strictly caudal direction, moving the platform by micromanipulator at the constant 5 mm/sec speed for approximately 10-15 mm. No or very little trunk movement was observed at stretching for the first 10 mm, and therefore the data recorded for this distance were included in statistical analysis. The force (in grams) from two gauges was simultaneously recoded with 100 Hz frequency during stretching. Five successive ramp-hold-return stretches were performed as technical replicates. Because the entire hindlimb was stretched the measured resistance was characteristic of the passive musculo-articular resistance integrated for hindlimb joints and muscles (Marsala et al., 2005; Nordez et al., 2008; Nordez et al., 2009). The resistance analyzed could have both neurogenic and mechanical components, but their respective contributions were not distinguished in the experimental design. The resistance was measured as the amount of mechanical work W_Left_ and W_Right_ to stretch the left- and right hindlimbs, where W was stretching force integrated over stretching distance interval from 0 to 10 mm.

## Statistical Analysis

Processing and statistical analysis of the HL-PA and stretching force data were performed after completion of the experiments by the statisticians, who were not involved in execution of experiments. Therefore, the results of intermediate statistical analyses could not affect acquisition of experimental data.

### Bayesian framework

Predictors and outcomes centered and scaled before we fitted Bayesian regression models via full Bayesian framework by calling *Stan* 2.21.2 from *R* 4.1.1 (Team, 2021) using the *brms* 2.16.1 (Burkner, 2017) interface. To reduce the influence of outliers, models used Student’s *t* response distribution with identity link function unless explicitly stated otherwise. Models had no intercepts with indexing approach to predictors (McElreath, 2020). Default priors were provided by the *brms* according to Stan recommendations (Gelman, 2019). Intercepts, residual SD and group-level SD were determined\from the weakly informative prior student_t(3, 0, 10). The additional parameter ν of Student’s distribution representing the degrees of freedom was obtained from the wide gamma prior gamma(2, 0.1). Group-level effects were determined from the very weak informative prior normal(0, 10). Four MCMC chains of 40000 iterations were simulated for each model, with a warm-up of 20000 runs to ensure that effective sample size for each estimated parameter exceeded 10000 (Kruschke, 2015) producing stable estimates of 95% highest posterior density continuous intervals (HPDCI). MCMC diagnostics were performed according to the Stan manual. P-values, adjusted using the multivariate t distribution with the same covariance structure as the estimates, were produced by frequentist summary in *emmeans* 1.6.3 (Searle et al., 2012) together with the medians of the posterior distribution and 95% HPDCI. The asymmetry and contrast between groups were defined as significant if the corresponding 95% HPDCI did not include zero and the adjusted P-value was ≤ 0.05. R scripts are available upon request.

### HL-PA. Postural asymmetry

The magnitude of postural asymmetry (MPA) was inferred via Bayesian framework using half-normal response distribution. The probability of HL-PA (P_A_) was inferred via Bayesian framework with Bernoulli response distribution and logit link function. For the asymmetric UBI rats, the probability of the contralesional flexion (P_C_) was modelled with binomial response distribution and logit link function. Significance of differences in P_C_ between the groups of asymmetric UBI rats and artificial “50/50” groups with equal number rats showing the left and right flexion used as a control, was computed.

### Stretching force

The amount of mechanical work W_Left_ and W_Right_ to stretch the left- and right hind limbs, respectively, was computed by integrating the smoothed stretching force measurements over stretching distance from 0 to 10 mm using loess smoothing computed by *loess* function from R package *stats* with parameters span=0.4 and family=“symmetric”. Asymmetry was assessed both as the left / right asymmetry index AI_LR_ = log_2_ (W_Left_ / W_Right_), and for contra- and ipsilesional hindlimbs AI_CI_ = log_2_ (W_Contra_ / W_Ipsi_) and as the difference in work, respectively ΔW_LR_ = (W_Left_ - W_Right_), ΔW_CI_ = (W_Contra_ – W_Ipsi_). The AI and ΔW were inferred via Bayesian framework by fitting linear multilevel models that included *operation type* (left UBI *vs*. right UBI *vs*. sham) as the factor of interest.

## Data availability

Data supporting the findings of this study are available within the article, its Supporting Information or upon request.

## Results

### The UBI-induced HL-PA in rats with completely transected cervical spinal cord

The UBI effects were studied in rats with the spinal cord transected at the cervical C6-C7 level. To ensure the completeness of the transection, a 3-4-mm spinal segment was excised in all analyzed rats, and then the hindlimb representation area of the sensorimotor cortex was unilaterally ablated. The UBI was performed by suction lesion in order to restrict a damaged area to the hindlimb sensorimotor cortex and examine specific changes in hindlimb motor functions. This lesion resulted in formation of HL-PA with contralesional hindlimb flexion in rats with intact brain along with contralesional hindlimb motor deficits in the beam-walking and ladder rung tests in rats with intact spinal cord as it was shown in previous experiments (Lukoyanov et al., 2021; Watanabe et al., 2021). HL-PA was analyzed before (designated as Pre) and 3 hours after the UBI or sham surgery (Post), by both the hands-on and hands-off methods of hindlimb stretching followed by photographic and / or visual recording of the asymmetry in animals under pentobarbital anesthesia (for details, see “Materials and methods”). Data obtained by these two methods well correlated (**Figure 1- Supplement 1**) and the results are presented for the hands-off assay (**Figures 1 and 2**). HL-PA was characterized by i) the postural asymmetry size (PAS) in mm (**Figures 1B,D and 2B,D**); ii) the magnitude of postural asymmetry (MPA) in mm (**Figure 2F,H**); and iii) the probability to develop HL-PA (P_A_) (**Figures 1C,E and 2C,E,G,I**). The rats with the MPA > 1 mm were defined as asymmetric; the 1 mm MPA was 94th percentile in rats before UBI or sham surgery and after sham surgery.

**Figure 1.**
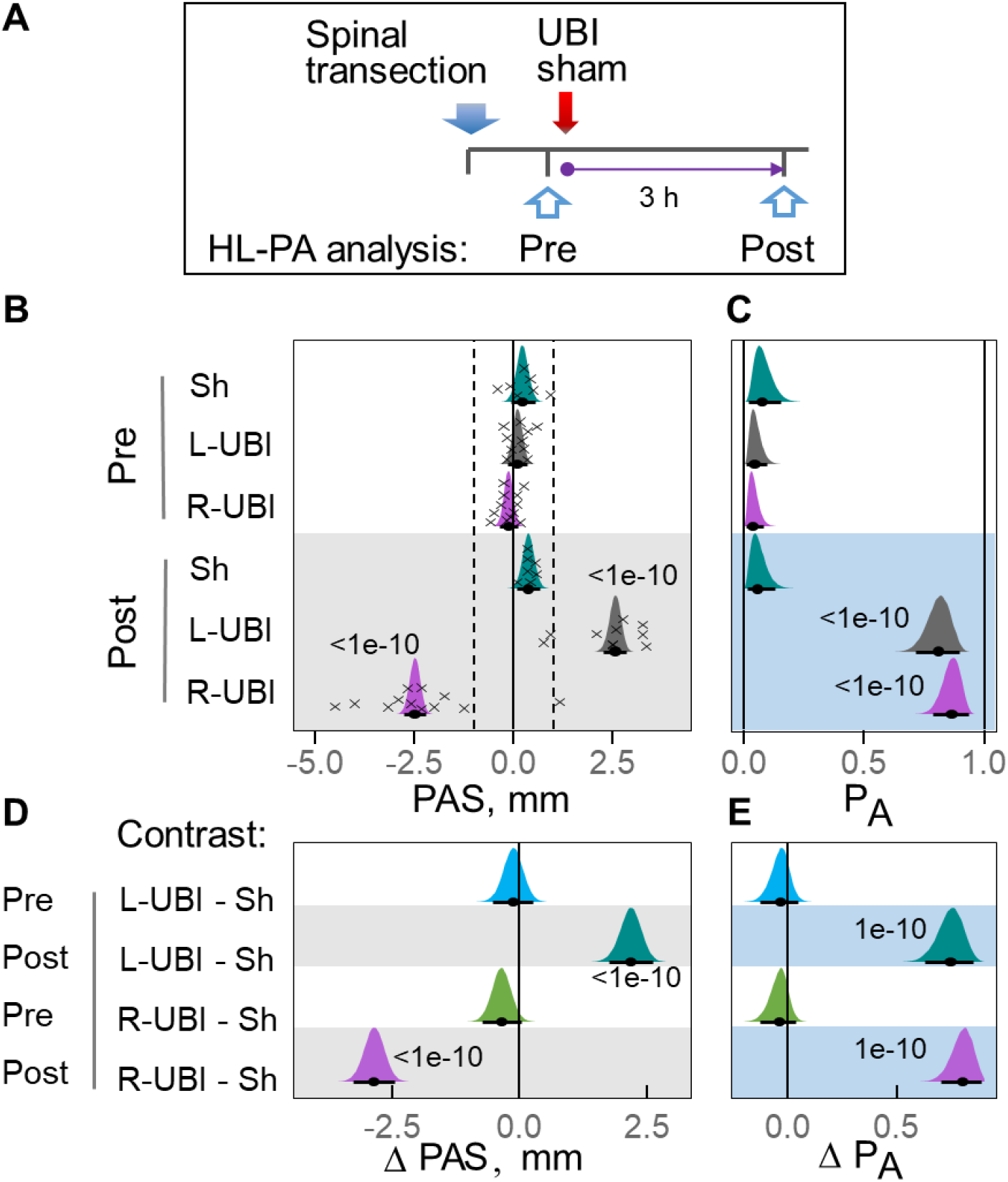
Postural asymmetry of hindlimbs (HL-PA) induced by the unilateral ablation of the hindlimb representation area of sensorimotor cortex (UBI) in rats with completely transected spinal cord. (**A**) Experimental design. The spinal cord was transected at the C6-7 level by the excision of 3 mm segment. HL-PA was analyzed before (Pre) and 3 h after (Post) left UBI (L-UBI; n = 10), right UBI (R-UBI; n = 12), or sham surgery (Sh; n = 7). Data of the hands-off analysis are shown. (**B**) The postural asymmetry size (PAS) in millimeters (mm), and (**C**) the probability to develop HL-PA (P_A_) above 1 mm threshold (denoted in **B** by vertical dotted lines). (**D,E**) Differences (contrasts) between the UBI and sham groups in the PAS and P_A_, before (Pre) and after (Post) UBI or sham surgery. The PAS, P_A_ and contrasts are plotted as median (black circles), 95% HPDC intervals (black lines), and posterior density (colored distribution) from Bayesian regression. Negative and positive PAS values are assigned to rats with the left and right hindlimb flexion, respectively. Significant effects on asymmetry and differences between the groups: 95% HPDC intervals did not include zero, and adjusted P-values were ≤ 0.05. Adjusted P is shown for differences identified by Bayesian regression. **Source data 1**. The EXCEL source data file contains data for panels (**B**) and (**C**) of ***Figure 1***. **Figure supplement 1**. High concordance of the hands-on and hands-off analyses of HL-PA.

**Figure 2.**
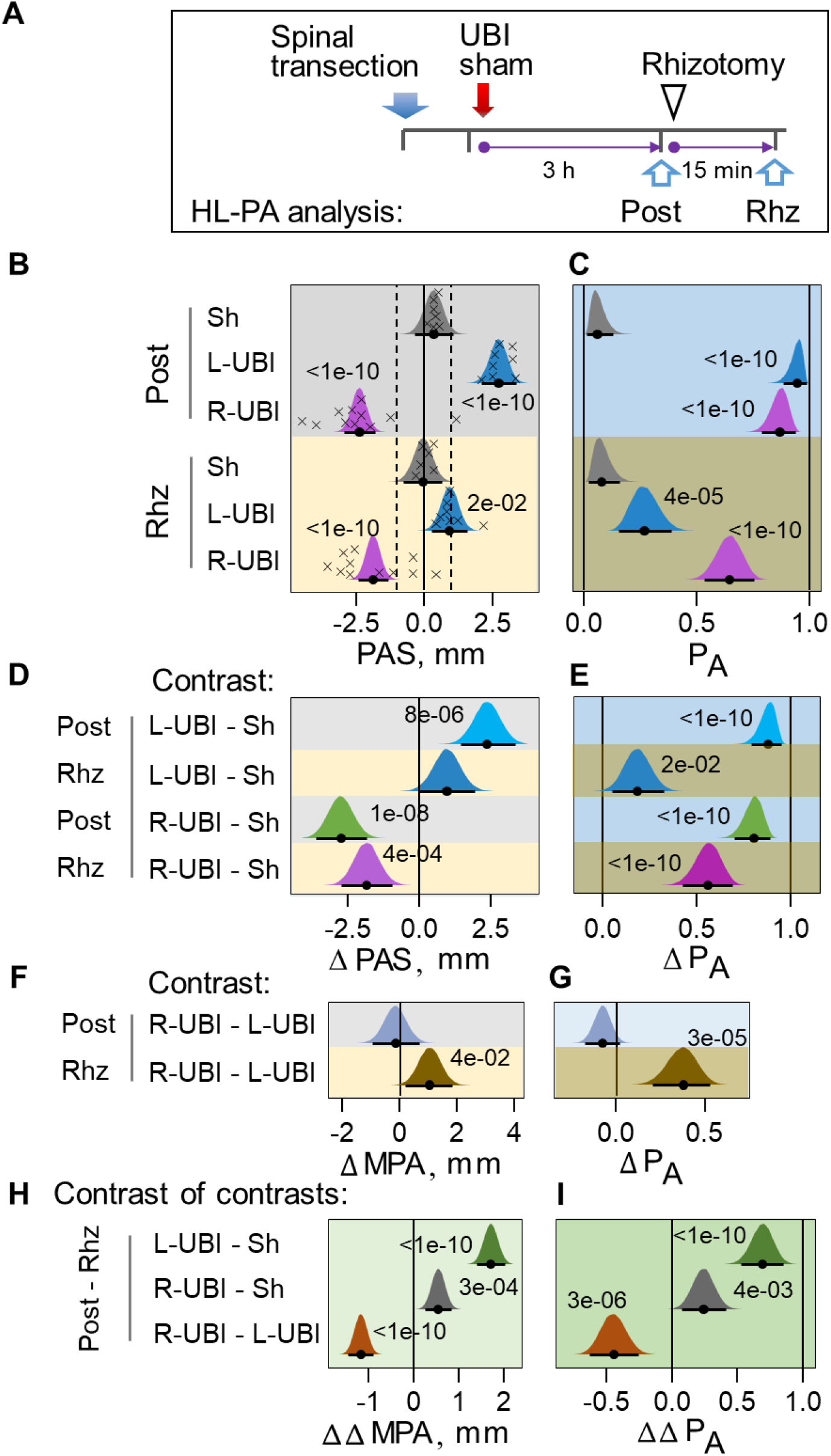
Effects of bilateral deafferentation by rhizotomy of lumbar spinal cord on formation of HL-PA induced by the L-UBI or R-UBI in rats with completely transected spinal cord. (**A**) Experimental design. The spinal cord was transected at the C6-7 level that was followed by L-UBI (n = 8), R-UBI (n = 11), or sham surgery (Sh; n = 7). Data of the hands-off analysis are shown. HL-PA was analyzed three hours after UBI or sham surgery (Post) and in the same rats after bilateral rhizotomy performed from the L1 to S2 spinal levels designated as (Rhz). (**B**) The postural asymmetry size (PAS) in millimeters (mm), and (**C**) the probability to develop HL-PA (P_A_) above 1 mm threshold (denoted in B by vertical dotted lines). (**D,E**) Differences (contrast) between the UBI and sham groups in the PAS and P_A_ for rat groups analyzed after UBI or sham surgery (Post) and the same rat groups after rhizotomy (Rhz). (**F,G**) Differences (contrasts) between the R-UBI and L-UBI groups in the magnitude of postural asymmetry (MPA) and P_A_ for the Post and Rhz rat groups. (**H,I**) The effects of rhizotomy on differences in the MPA (or P_A_) between L-UBI, R-UBI and sham surgery (Sh) were analyzed as contrast of contrasts i) between L-UBI and sham surgery: Δ ΔMPA (or Δ ΔP_A_) = [(L-UBI _Post_ – Sh _Post_) – [(L-UBI _Rhz_ – Sh _Rhz_)]; ii) between R-UBI and sham surgery: Δ ΔMPA (or Δ ΔP_A_) = [(R-UBI _Post_ – Sh _Post_) – [(R-UBI _Rhz_ – Sh _Rhz_)]; and iii) between R-UBI and L-UBI sham surgery: Δ ΔMPA (or Δ ΔP_A_) = [(R-UBI _Post_ – L-UBI _Post_) – [(R-UBI _Rhz_ – L-UBI _Rhz_)]. The PAS, P_A_, MPA and contrasts are plotted as median (black circles), 95% HPDC intervals (black lines), and posterior density (colored distribution) from Bayesian regression. Negative and positive PAS values are assigned to rats with the left and right hindlimb flexion, respectively. Effects on asymmetry and differences between the groups: 95% HPDC intervals did not include zero, and adjusted P-values were ≤ 0.05. Adjusted P is shown for differences identified by Bayesian regression.

In rats with transected cervical spinal cords, the UBI induced HL-PA (**Figure 1B-E**). The PAS and the probability of postural asymmetry in rats with left (n = 10) and right (n = 12) UBI were much greater, 7-fold or more than in rats before UBI or sham surgery (n = 7), or after sham surgery. The probability and the MPA did not differ between the left- and right-side UBI groups (**Figures 1C,E** and **2F, G**). Differences (contrast) in the PAS and PA between the UBI and sham surgery groups were highly significant (**Figure 1D, E**). Strikingly, the response to the injury developed on the contralesional side; the left or right UBI induced the right and left hindlimb flexion, respectively (**Figure 1B,D**). The PAS, the probability, and formation of contralesional hindlimb flexion induced by UBI in rats with transected cervical spinal cords were similar to the previously reported respected values for UBI animals with intact spinal cords (Lukoyanov et al., 2021). We conclude that HL-PA formation in animals with transected cervical spinal cords is not mediated by the descending neural tracts and the sympathetic system, and that an extraspinal, likely humoral mechanism conveys a message on the brain injury and its side from the brain to the lumbar spinal cord.

### Effects of bilateral deafferentation of lumbar spinal segments on HL-PA induced by the L-UBI or R-UBI

We next sought to determine whether afferent somatosensory input is required for the HL-PA formation in the UBI rats with transected spinal cord. The HL-PA was analyzed before and after bilateral rhizotomy from the L1 to S2 spinal levels that was performed 3 hours after injury either of the left or right hemisphere or sham surgery in rats with transected cervical spinal cord. In rats with left UBI, the PAS and P_A_ were strongly, 3.0- and 3.5-fold, respectively reduced after rhizotomy (**Figure 2**). In contrast, in the right-side UBI rats, the PAS and P_A_ demonstrated small, approximately 1.3-fold decrease after rhizotomy. No effects of rhizotomy were evident in rats with sham surgery.

While before rhizotomy contrast between both the UBI groups and sham surgery group in each the PAS and PA was strong and highly significant, after it there was no or weak difference in the left UBI group (**Figure 2D,E**). The right UBI – sham surgery group contrast remained strong and significant after rhizotomy. No contrast between the left and right UBI groups was evident before rhizotomy while after it the differences was significant (**Figure 2F,G**). Relative impact of rhizotomy (before vs. after it) on the effects of left and right UBI (both UBI vs. sham surgery) was analyzed as contrast of contrasts (**Figure 2H,I**). In other words, contrast in each the MPA and the probability of postural asymmetry between the animal groups (left UBI vs. sham surgery; right UBI vs. sham surgery; and left UBI vs. right UBI) was compared between two time points that were before vs. after rhizotomy. While contrast with sham group was high and significant for the left UBI group, it was much smaller for the right UBI rats. Contrast of contrasts for the left UBI vs. right UBI was also strong and significant.

These data suggest that the HL-PA formation in rats with transected spinal cord depend on i) afferent somatosensory input, and ii) activity of motoneurons signaling to hindlimb muscles after the left-side and right-side UBI, respectively. Reversibility of the HL-PA after left UBI by deafferentation also suggests that neuroplastic changes in the lumbar neurocircuits could not account for the asymmetry formation. In contrast, the neuroendocrine signals after the right side UBI may induce formation of a pathological trace in efferent neurocircuits that remain active after rhizotomy in the lumbar spinal cord.

### Hindlimb stretching resistance: effects of UBI

We next studied the UBI effects on biomechanical properties of the contra- and ipsilesional hindlimbs. The passive hindlimb musculo-articular resistance to stretching were assessed in the anesthetized rats with transected cervical spinal cords 3 hours after UBI or sham surgery (**Figures 3,4 and Figure 4-supplement 1**). The resistance force was analyzed using the micromanipulator-controlled force meter device that consisted of two digital force gauges fixed on a movable platform (**Figure 3A**). Two silk threads were glued to the hindlimb nails of rats placed in symmetric position. Their other ends were hooked to the force gauges, and then the hindlimbs were stretched. The asymmetry in the resistance was assessed as i) the difference in the work (ΔW) between the left- and right hindlimbs ΔW_LR_ = (W_Left_ – W_Right_) where W_Left_ and W_Right_ were the W applied to stretch the left- and right hindlimbs (**Figure 4B-E**); and ii) the left / right asymmetry index for the work, AI_LR_ = log_2_ (W_Left_ / W_Right_) (**Figure 4-supplement 1B-E**). The effects of left and right UBI were compared by computing the indexes ΔW_CI_ = (W_C_– W_I_) (**Figure 4F**) and AI_CI_ = log_2_ (W_C_ / W_I_) (**Figure 4-supplement 1F**) for difference in the work between contra- (C) and ipsilesional (I) hindlimbs. The indexes were integrated for a 10-mm stretching distance. Both the W and AI were analyzed because they may differently depend on the stretching distance.

**Figure 3.**
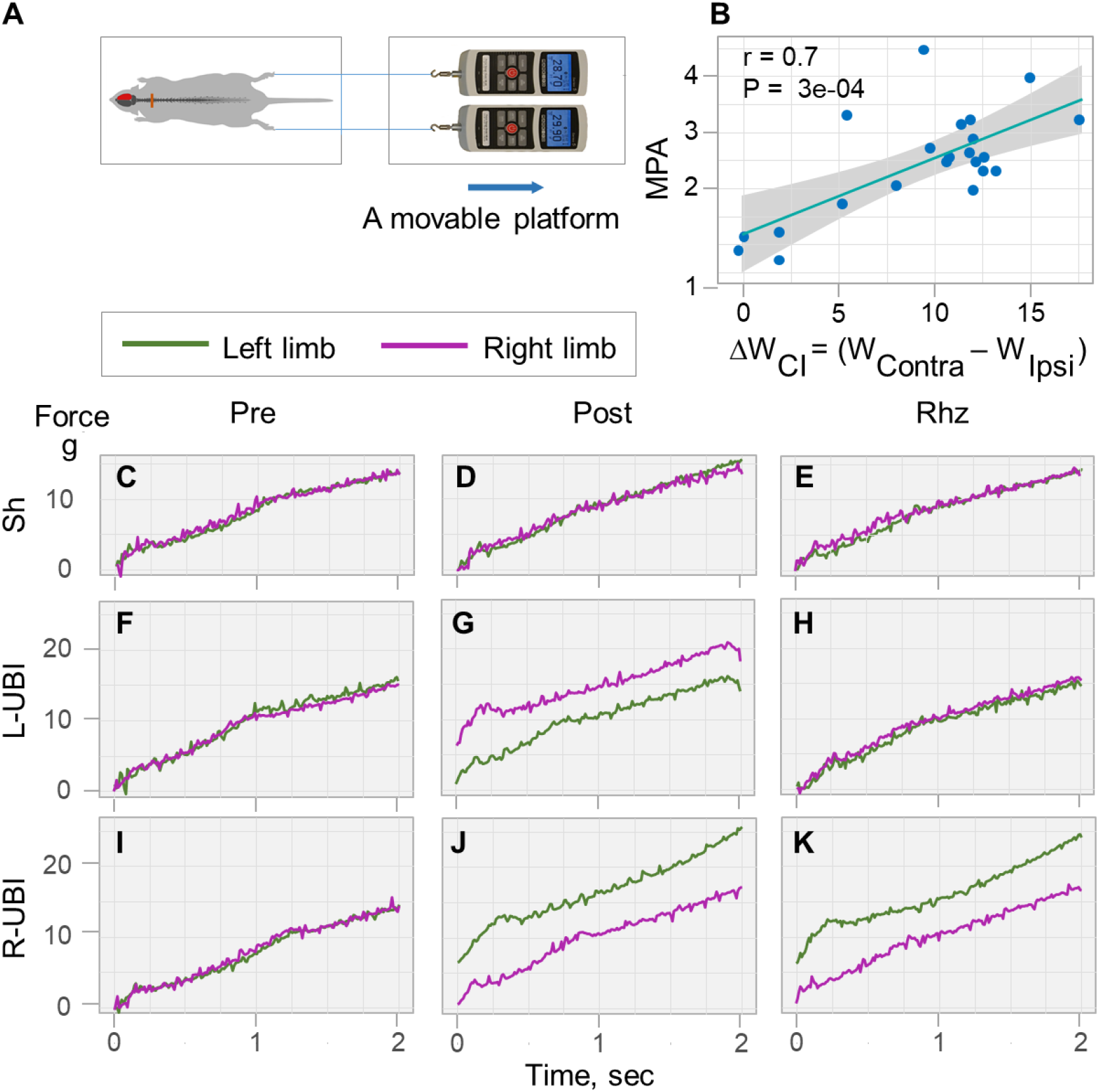
Analysis of the stretching force of the left and right hindlimbs in rats with transected spinal cord. Representative traces were recorded before and after UBI or sham surgery, and then after rhizotomy. (**A**) Experimental setup. The micromanipulator-controlled force meter device consisted of two digital force gauges fixed on a movable platform operated by a micromanipulator. The rat was placed on a heating pad. Two silk threads were hooked to the force gauges and their other ends were glued to the nails of the middle three toes of each hindlimb. The legs were placed in symmetric position and stretched for 10 mm distance at 5 mm/sec speed. The stretching resistance was analyzed as the amount of mechanical work W to stretch a hindlimb, calculated as integral of stretching force over 0-10 mm distance. (**B**) Pearson correlation between the postural asymmetry magnitude (MPA) analyzed by the hands-on assay and differences in the work between the contra- and ipsilesional hindlimbs [ΔW_CI_ = (W_Contra_ – W_Ipsi_)] in gm × mm. Data are presented for L-UBI and R-UBI groups analyzed 3 hours after brain surgery. (**C-K**) Representative traces of the stretching force recorded from the left and right hindlimbs of rats with spinal cord transected at the C6-7 level before UBI and sham operation (Pre), 3 h after brain surgery (Post), and then after rhizotomy (Rhz).

**Figure 4.**
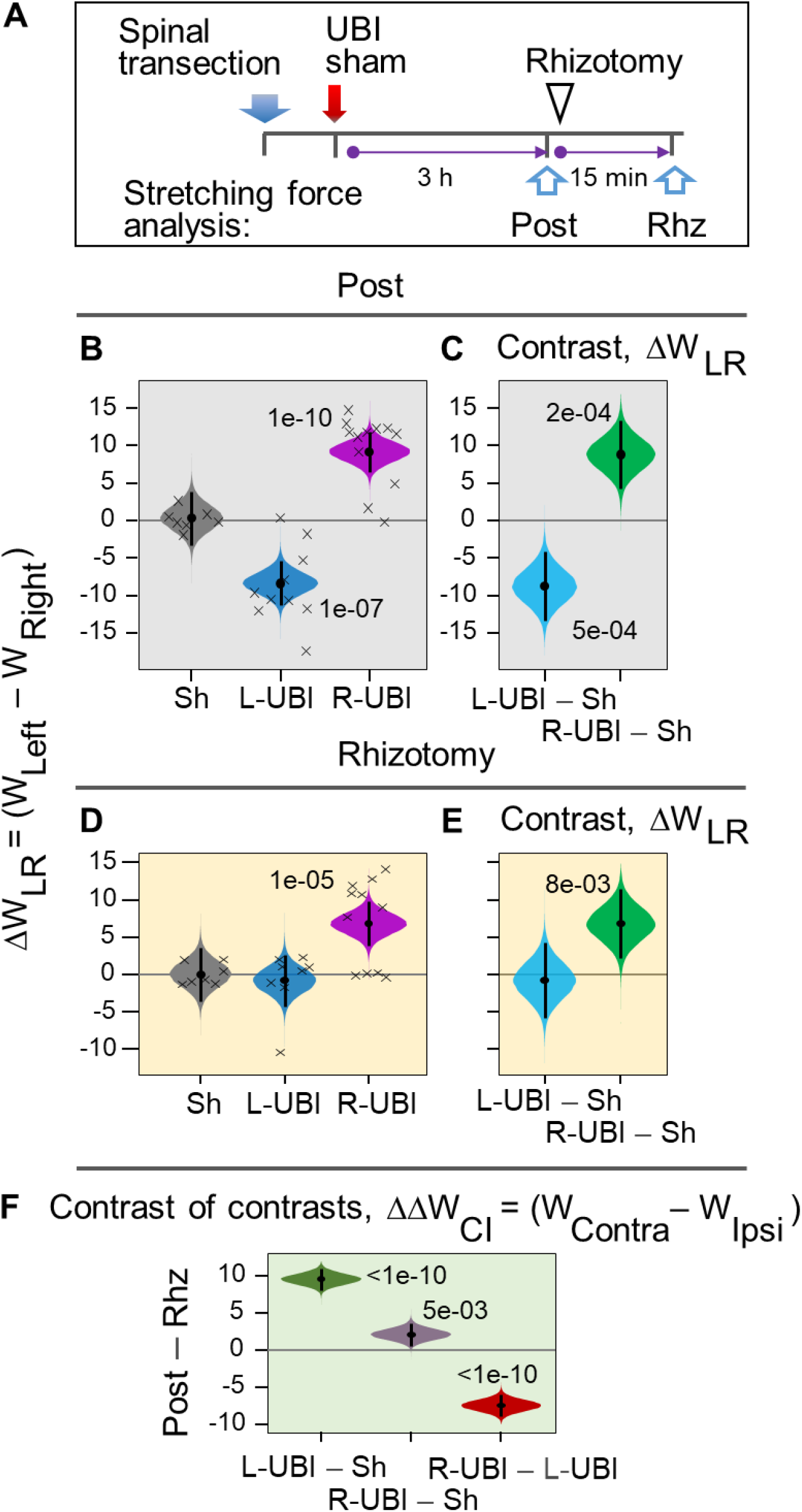
Stretching resistance of the left and right hindlimbs in rats with completely transected spinal cord: effects of UBI and bilateral deafferentation by rhizotomy of lumbar spinal cord. (**A**) Experimental design. The spinal cord was transected that was followed by L-UBI, R-UBI, or sham surgery (Sh). Stretching force was analyzed (**B,C**) three hours after UBI or sham surgery (Post; L-UBI, n = 10; R-UBI, n = 12; and sham surgery, n = 7); and (**D,E**) after bilateral rhizotomy designated as Rhz in the subset of rats (L-UBI, n = 8; R-UBI, n = 11; and sham surgery, n = 7). Differences in stretching force were calculated between the left and right hindlimbs as ΔW_LR_ = (W_Left_ - W_Right_) in (**B**-**E**), or between the contra and ipsilesional hindlimbs as ΔW_CI_ = (W_Contra_ – W_Ipsi_) in (**F**) in gm × mm. (**C,E**) Differences (contrast) between the UBI and sham surgery groups. (**F**) The effects of rhizotomy on differences in ΔW_CI_ between L-UBI, R-UBI and sham surgery were analyzed as contrast of contrasts Δ Δ i) between L-UBI and sham surgery: [(L-UBI _Post_ – Sh _Post_) – [(L-UBI _Rhz_ – Sh _Rhz_)]; ii) between R-UBI and sham surgery: [(R-UBI _Post_ – Sh _Post_) – [(R-UBI _Rhz_ – Sh _Rhz_)]; and iii) between R-UBI and L-UBI sham surgery: [(R-UBI _Post_ – L-UBI _Post_) – [(R-UBI _Rhz_ – L-UBI _Rhz_)]. The W_LR_, W_CL_, contrast and contrast of contrasts are plotted as median (black circles), 95% HPDC intervals (black lines), and posterior density (colored distribution) from Bayesian regression. Significant effects on the ΔW and the differences between the groups: 95% HPDC intervals did not include zero, and adjusted P-values were ≤ 0.05. Adjusted P is shown for differences identified by Bayesian regression.

Representative traces of the stretching force recorded from left and right hindlimbs of rats with transected cervical spinal cord before and 3 h after UBI or sham operation are shown on **Figure 3C-K**. Force to stretch hindlimbs increased during stretching starting from zero. No differences were evident in the stretching force between the contra and ipsilesional hindlimbs before sham surgery and UBI and after sham surgery (**Figure 3C,D,F,I**). The left or right UBI induced marked increase in the stretching resistance of the contralesional limb that was analyzed 3 hours after brain operation (**Figure 3G, J**). This pattern was observed at all-time points during stretching. The difference in the work between the contra and ipsilesional limbs (ΔW) strongly correlated with formation of HL-PA (the ΔW *vs*. the MPA: r = 0.7; P = 3e-04) (**Figure 3B**).

Differences in the work applied to stretch the left and right hindlimbs (ΔW) and in the AI were highly significant after left and right UBI while sham surgery did not produce the asymmetry (**Figure 4B; Figure 4-figure supplement 1B**). The contralesional hindlimbs demonstrated markedly higher resistance in comparison to ipsilesional hindlimbs. Both the left and right UBI groups significantly differed from the sham group in the ΔW (**Figure 4C**) and the AI (**Figure 4- figure supplement 1C**). Thus in rats with completely transected cervical spinal cord each the left and right UBI produced robust and highly significant increase in the musculo-articular resistance to stretching of the contralesional vs. ipsilesional hindlimbs.

### Effects of bilateral deafferentation on the UBI-induced asymmetry in hindlimb stretching resistance

The stretching resistance of the contra- and ipsilesional hindlimbs in rats with transected cervical spinal cords was analyzed before and after bilateral rhizotomy from the L1 to S2 spinal levels that was performed 3 hours after UBI or sham surgery (**Figure 4D-F; Figure 4-figure supplement 1D-F**). In rats with left UBI rhizotomy abolished the differences between the hindlimbs in the resistance (**Figures 3H and 4D; Figure 4-figure supplement 1D**). In contrast, rats with the right UBI still displayed strong and significant asymmetry after rhizotomy (**Figures 3K and 4D; Figure 4-figure supplement 1D**). No rhizotomy effects were evident in rats with sham surgery (**Figures 3E**). While before rhizotomy contrasts the ΔW_LR_ and the ΔAI_LR_ between each UBI group and sham surgery group were strong and highly significant, after it there were virtually no differences in the left UBI group (**Figure 4E; Figure 4-figure supplement 1E**). Contrasts between the right UBI – sham surgery groups remained strong and significant after rhizotomy.

Impact of rhizotomy (contrast: before vs. after it) on the effects of left and right UBI (contrast: UBI vs. sham surgery) was analyzed as contrast of contrasts (**Figure 4F; Figure 4-figure supplement 1F**). Namely, contrast in the ΔW_CI_ and the ΔAI_CI_ (left UBI vs. sham surgery; right UBI vs. sham surgery; and left UBI vs. right UBI) was compared between two measurements that were performed before and after rhizotomy. This analysis revealed that in the left UBI group contrast with sham group was high and significant, while it was markedly smaller for the right UBI rats. Contrast of contrasts for the left UBI vs. right UBI was strong and highly significant. These data may be interpreted as strong and significant effects of deafferentation on the asymmetric effects induced by the left UBI, while negligible changes were induced by deafferentation in rats with the right UBI. These findings corroborate results of HL-PA analysis and suggest that the effects of the left-side and right-side UBI in rats with transected spinal cord depend on i) afferent somatosensory input, and ii) activity of motoneurons signaling to hindlimb muscles, respectively. Different effects of the left and right-side rhizotomy on interneuronal circuitry that lead to different motor outputs for the left and right limbs could not be ruled out.

## Discussion

### The left-right side-specific humoral signaling

Our previous (Lukoyanov et al., 2021) and present studies provide evidence that the left-right side-specific humoral signaling mediates the effects of UBI on the formation of HL-PA, asymmetry in passive hindlimb musculo-articular resistance to stretching and withdrawal reflexes, and asymmetric changes in gene expression patterns in the lumbar spinal cord. Hypothetically, the paravertebral chain of sympathetic ganglia, which is the remaining neural connection between the rostral and caudal parts of the transected spinal cord, may convey supraspinal signals to the muscle vasculature and through this mechanism may differentially affect ipsi- and contralesional muscles. The following findings question this hypothesis. Sympathetic ganglia do not mediate control of lumbar neural circuits by the supraspinal structures (Brodal, 1981; Wolpaw and Lee, 1989), and the sympathetic system has a limited capacity to independently regulate blood flow to the left and right hindlimbs (Lee et al., 2007). A critical evidence that the sympathetic system does not mediate signals from injured brain to the lumbar spinal cord is provide by the present study; the hindlimb postural asymmetry and asymmetry in their musculo-articular resistance to stretch develop in rats with complete transection of the cervical spinal cord that was at the level rostral to the localization of preganglionic sympathetic neurons in the T1 - L2/3 segments. Experiments with hypophysectomized rats, “pathological” serum and neurohormones that induce HL-PA provide evidence for the humoral pathway in rats with transected spinal cord (Lukoyanov et al., 2021).

### Left-right side-specific targeting of the afferent vs. efferent spinal circuits by humoral signaling

HL-PA and asymmetry in hindlimb resistance to stretching induced by the right-side UBI are not affected by bilateral deafferentation of the lumbar spinal cord suggesting that spinal reflexes do not contribute to asymmetry formation. In contrast, bilateral lumbar rhizotomy after the left-side UBI abolishes HL-PA that thereby may develop due to differences in reflexes between the contra- and ipsilesional sides. Thus, the left-right side-specific humoral mechanism in animals with transected spinal cords may act through afferent somatosensory input after the left-side UBI, and, in contrast, by activation of motoneurons signaling to hindlimb muscles or changes in neuromuscular system after the right-side injury.

Neurological phenomena are often developed differently after injury to the left and right hemisphere. The right-side compared to the left-side stroke results in poorer postural responses in quiet and perturbed balance suggesting a prominent role of the right hemisphere in efferent control of balance (Fernandes et al., 2018). Hemineglect and bias of the subjective vertical are general causes of postural asymmetry and instability (Tasseel-Ponche et al., 2015). The serious postural impairment - the contraversive pushing called “Pusher syndrome”, which is linked to biased verticality and hemispatial neglect, is more common among patients with the right rather than left-hemisphere lesions (Perennou et al., 2008). Components in trunk control may be impaired depending on the side of the lesion. “Postural instability” is significantly more frequent among patients with right-hemisphere lesions, while “apraxic responses” predominate among those with left-hemisphere injury (Spinazzola et al., 2003). HL-PA that uncovers the left-right side-specific effects of brain lesions may be a convenient animal model to experimentally address the aforementioned clinical features.

Clinical studies demonstrated that a large fraction of subjects with stroke and cerebral palsy do not relax their muscles – they are tonically constricted without any voluntary command. This phenomenon is called spastic dystonia and defined as “stretch- and effort-unrelated sustained involuntary muscle activity following central motor lesions” (Gracies, 2005; Lorentzen et al., 2018). This clinical phenomenon can alter posture at rest thereby contributing to the hemiplegia (Marinelli et al., 2017) and is regarded as a form of efferent muscle hyperactivity (Baude et al., 2018; Gracies, 2005). Spastic dystonia may have a central mechanism which does not depend on afferent input in contrast to spasticity based on exacerbated reflex excitability (Sheean and McGuire, 2009). It was postulated that rhizotomy which abolished sensory input and diminish stretch reflexes would have no effect on spastic dystonia. The example is the absence of effects of dorsal rhizotomy on development of muscle contractures in children with cerebral palsy (Tedroff et al., 2015). Spastic dystonia was reported in monkeys with lesions of motor cortex that displayed involuntary muscle activity (Denny-Brown, 1966; Denny-Brown, 1980). The dystonia persisted after disruption of sensory input to the spinal cord suggesting that a central mechanism but not exaggerated reflexes were involved. Our findings suggest that spastic dystonia may develop more frequently in the left hindlimb after injury to the right hemisphere than in that on the right side.

### Translation of humoral messages into left-right side-specific responses

Besides encoding of information about the injury side into humoral messages, their translation into the left-right side specific responses at peripheral nerve endings or spinal neurons, is the key stage of the revealed phenomenon. We identified several neurohormones including opioid peptides and Arg-vasopressin, and synthetic agonists of selective opioid receptors that induced HL-PA after their intrathecal or intravenous administration into rats with intact brain (Bakalkin et al., 1981; Bakalkin and Kobylyansky, 1989; Bakalkin et al., 1986; Chazov et al., 1981; Lukoyanov et al., 2021; Watanabe et al., 2020). The critical finding is that the side of the flexed limb depends on the compound administered. κ-Opioid agonists dynorphin, bremazocine, and U-50,488, along with the μ/δ-opioid agonist Met-enkephalin induce flexion of the left hindlimb (Bakalkin et al., 1981; Bakalkin and Kobylyansky, 1989; Bakalkin et al., 1986; Chazov et al., 1981; Watanabe et al., 2020). In contrast, β-endorphin and Arg-vasopressin, and the δ-agonist Leu-enkephalin, cause the right limb to flex (Bakalkin et al., 1981; Chazov et al., 1981; Klement’ev et al., 1986; Lukoyanov et al., 2021).

Topographical information on the brain injury and its side may be conveyed by these molecular messengers released into the blood and then converted into side-specific motor responses at their target sites. Consistently an excision of the pituitary gland, the main source of these neurohormones in the circulation disables the humoral pathway and abolishes the HL-PA formation. Furthermore, serum from UBI rats induces HL-PA with contralesional hindlimb flexion in rats with intact brain. Each SSR-149415, a selective antagonist of the V1B vasopressin receptors mostly expressed in the pituitary gland, and naloxone, a non-selective opioid antagonist blocks the left UBI-induced formation of HL-PA in rats with transected spinal cords. These findings demonstrate that Arg-vasopressin and β-endorphin may transmit signals from the brain injured on the left side to the spinal neurocircuits controlling the right hindlimb functions.

A number of studies revealed left–right asymmetry in the spinal cord organization (de Kovel et al., 2017; Deliagina et al., 2000; Hultborn and Malmsten, 1983a; Hultborn and Malmsten, 1983b; Kononenko et al., 2017; Malmsten, 1983; Nathan et al., 1990; Ocklenburg et al., 2017; Zhang et al., 2020). Three-quarters of cervical spinal cords are asymmetric with a larger right side (Nathan et al., 1990). Mono- and polysynaptic segmental reflexes evoked by stimulation of the dorsal roots and recorded in the ventral roots in intact rats and cats display higher activity on the right side (Hultborn and Malmsten, 1983a; Hultborn and Malmsten, 1983b; Malmsten, 1983). Similarly, hindlimb withdrawal reflexes evoked by electrical stimulation display higher activity on the right side (Zhang et al., 2020). These asymmetric neural circuits controlling the left and right limbs may be targeted by opioid peptides and Arg-vasopressin released into the blood after the UBI (Bakalkin et al., 1981; Bakalkin et al., 1986; Chazov et al., 1981; Klement’ev et al., 1986). The left-right side specific neurohormonal effects may be mediated through their receptors lateralized in the spinal cord. Indeed, in the rat lumbar segments, the expression of three opioid receptors is lateralized to the left, and their proportions and co-expression patterns are different between the left and right sides (Kononenko et al., 2017; Watanabe et al., 2021). This asymmetry may enables translation of the hormonal messages into the left-right side specific response after a unilateral brain injury.

## Limitations

The side-specific humoral signaling was revealed in anaesthetized animals with transected spinal cords that were studied during 3 – 5 hours after UBI. These experiments did not allow to identify biological and pathophysiological relevance of this phenomenon. Pathways from the injured brain area to the hypothalamic-pituitary system, neurohormones mediating effects of the right-side injury, peripheral or central targets for the left- and right-side specific endocrine messengers, and afferent, central or efferent mechanisms of the asymmetry formation have not been investigated.

This study does not focus on mechanisms and clinical correlates of postural and sensorimotor deficits. Hindlimb posture and resistance to stretch were studied as readouts of the UBI because they are regulated by neurohormones and may be analyzed after spinal cord transection. Furthermore, they are directional – polarized along the left-right axis, and, therefore, can be used to examine if the humoral pathway conveys the side-specific signals. At the same time, both the HL-PA and the resistance model several features of the brain injury-induced sensorimotor and postural deficits in humans. First, the changes induced by UBI in hindlimb functions have a contra-ipsilesional pattern. Second, the HL-PA after the right-side injury does not depend on the afferent input and in this regard it may be similar to “spastic dystonia”, a tonic muscle overactivity that contributes to “hemiplegic posture” (Gracies, 2005; Lorentzen et al., 2018; Sheean and McGuire, 2009). Third, asymmetric exacerbated withdrawal reflexes that lead to flexor spasms in patients (Bussel et al., 1989; Dietz et al., 2009; Lavrov et al., 2006; Schouenborg, 2002) are similarly developed in rats (Lukoyanov et al., 2021; Zhang et al., 2020).

## Conclusion

Functional specialization of the left and right hemispheres is an organizing principle of the brain (Concha et al., 2012; Duboc et al., 2015; MacNeilage et al., 2009; Ocklenburg et al., 2017). The computational advantages of the specialization may include paralleled processing of information modules in the left and right hemispheres that increases the flow of information, and improves functional performance. Lasting regulation of the lateralized processes may be accomplished by local paracrine mechanisms that differentially operate on the left and right side of the midline through released neurohormones (Allen et al., 2021; Hussain et al., 2012; Kononenko et al., 2017; Kononenko et al., 2018; Marlin et al., 2015; Nation et al., 2018; Phelps et al., 2019; Watanabe et al., 2015; Watanabe et al., 2020; Zink et al., 2011). Our findings suggest a more general role for the lateralized neuropeptide systems than regulation of the lateralized functions. We hypothesize that the left- and right-side specific neurohormonal mechanism regulates the mirror-symmetric neural circuits that control paired organs and extremities including the left and right hindlimbs. Neurohormones may differentially target the left and right counterparts of these circuits, and by this virtue control left-right balance in their functional performance. This bipartite mechanism may be based on lateralization of the neurohormonal systems, and may operate locally (e.g., within the lumbar spinal cord), or at the several levels of the neuraxis by controlling neural pathways that convey signals from the left and right hemispheres to the contralateral hindlimb motoneurons. Unilateral brain or body lesion may shift this balance to the left or to the right, depending on the side of injury, and thereby impair the left-right side specific neurohormonal control leading to asymmetric functional deficits.

In conclusion, this and previous works present evidence for the novel phenomenon, the side-specific humoral mechanism that mediates asymmetric effects of unilateral brain injury on hindlimb posture and sensorimotor functions. The humoral pathway and the descending neural tracts may represent complementary routes for signaling from the brain to the spinal cord. Analysis of features and proportion of postural and sensorimotor deficits transmitted by neurohormonal signals versus those mediated by neural pathways in animal models and patients after stroke and traumatic brain injury should facilitate new therapeutic discoveries. From a biological standpoint, the mechanism may serve to maintain a balance between the left–right processes in bilaterally symmetric animals.

## Author contribution

H.W. and M.Z., performed injury, behavioral and morphological analysis. Y.K. and D.S. performed statistical analyses. H.W., M.Z., I.L., J.S. and G.B. discussed the data and participated in manuscript preparation. J.S., M.Z. and G.B. conceived and supervised the project. G.B. wrote the manuscript. All authors worked with and commented on the manuscript.

## Acknowledgements

We are grateful to Dr. Michael Ossipov for discussion and manuscript processing.

