## Supplemental Data 1 for "Humoral signaling-mediated effects of unilateral brain injury: differences in the left-right sided afferent responses"

**Supplementary figures and figure legends**

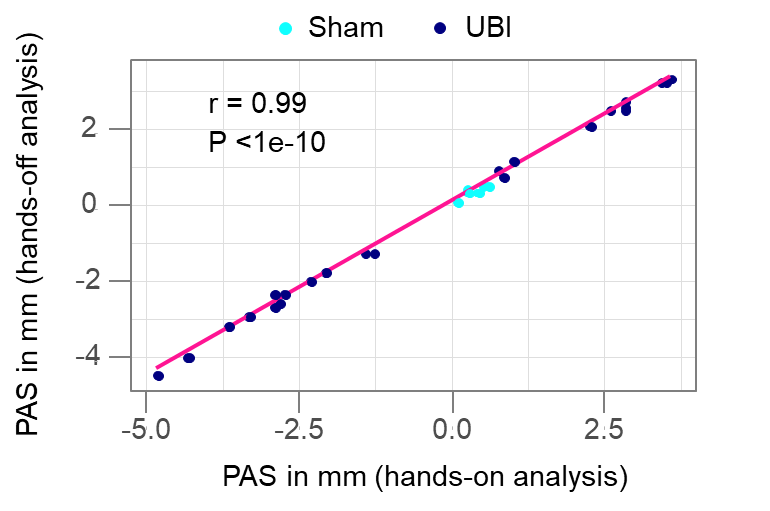

**Figure 1—figure supplement 1. Pearson correlation between the postural asymmetry size (PAS) analyzed by the hands-off and hands-on assay.** Data were combined for L-UBI, R-UBI and sham surgery groups of rats with transected spinal cord that were analyzed 3 hours after brain surgery.

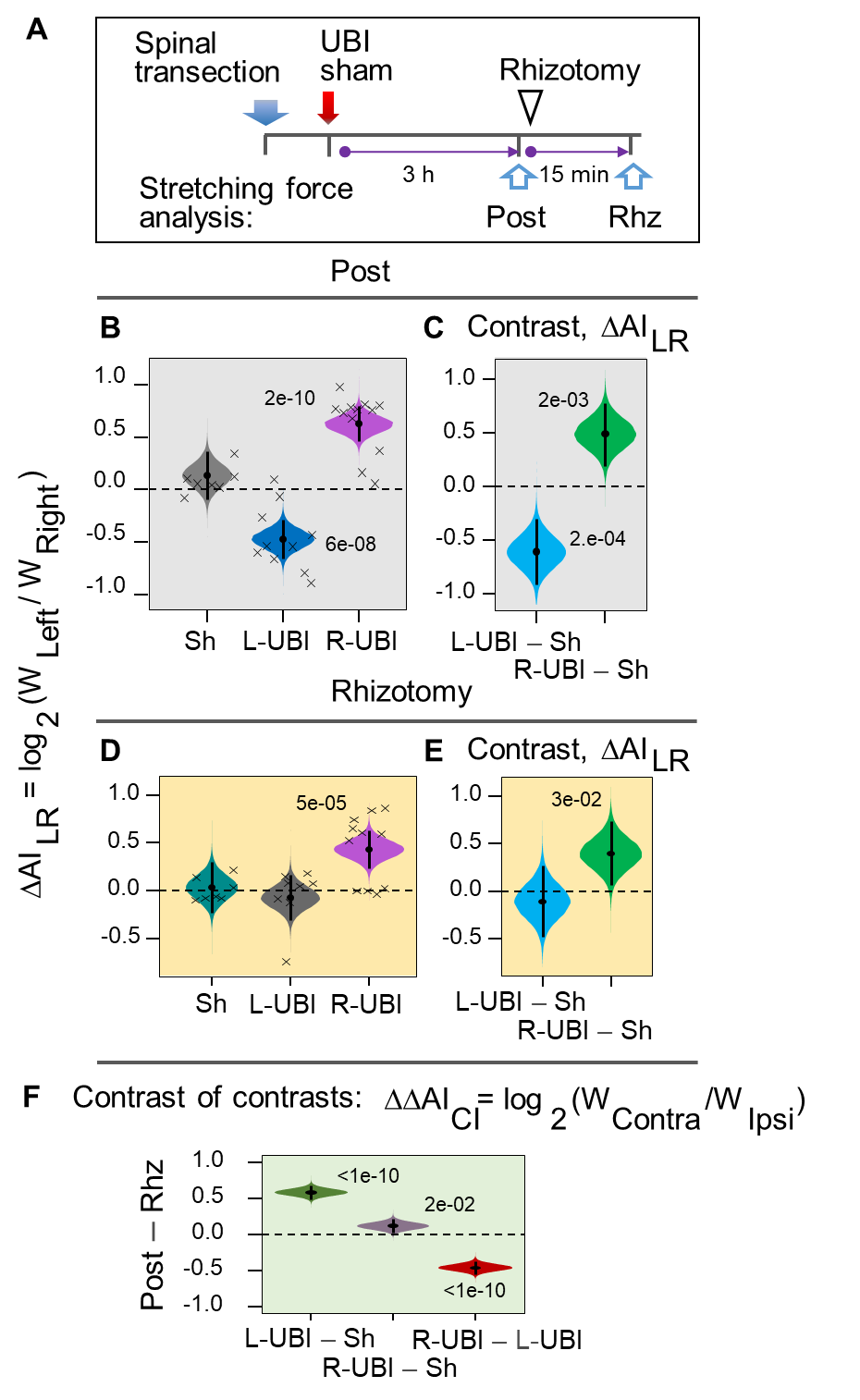

**Figure 4_Supplement 1. Asymmetry in the hindlimb stretching resistance in rats with completely transected spinal cord: effects of UBI and bilateral deafferentation by rhizotomy of lumbar spinal cord.** (**A**) Experimental design. The spinal cord was transected that was followed by L-UBI, R-UBI or sham surgery (Sh). Stretching force was analyzed (**B**,**C**) three hours after UBI or sham surgery (Post; L-UBI, n = 10; R-UBI, n = 12; and sham surgery, n = 7); and (**D**,**E**) after bilateral rhizotomy designated as Rhz in the subset of rats (L-UBI, n = 8; R-UBI, n = 11; and sham surgery, n = 7). The UBI effects were analyzed as changes in the asymmetry index for left and right hindlimbs AI_LR_ = log_2_ (W_Left_ / W_Right_) (**B-E**), and for contra- and ipsilesional hindlimbs AI_CI_ = log_2_ (W_Contra_ / W_Ipsi_). (**C**,**E**) Differences (contrast) between the UBI and sham surgery groups. (**F**) The effects of rhizotomy on differences in AI_CI_ between L-UBI, R-UBI and sham surgery were analyzed as contrast of contrasts i) between L-UBI and sham surgery: [(L-UBI _Post_ – Sh _Post_) – [(L-UBI _Rhz_ – Sh _Rhz_)]; ii) between R-UBI and sham surgery: [(R-UBI _Post_ – Sh _Post_) – [(R-UBI _Rhz_ – Sh _Rhz_)]; and iii) between R-UBI and L-UBI sham surgery: [(R-UBI _Post_ – L-UBI _Post_) – [(R-UBI _Rhz_ – L-UBI _Rhz_)]. The AI_LR_, AI_CL_, contrast and contrast of contrasts are plotted as median (black circles), 95% HPDC intervals (black lines), and posterior density (colored distribution) from Bayesian regression. Significant effects on the ΔW and the differences between the groups: 95% HPDC intervals did not include zero, and adjusted P-values were ≤ 0.05. Adjusted P is shown for differences identified by Bayesian regression.
